# genuMet: distinguish genuine untargeted metabolic features without quality control samples

**DOI:** 10.1101/837260

**Authors:** L Cao, C Clish, FB Hu, MA Martínez-González, C Razquin, M Bullo-Bonet, D Corella, E Gómez-Gracia, M Fiol, R Estruch, J Lapetra, M Fitó, F Arós, L Serra-Majem, E Ros, L Liang

## Abstract

**Motivation:** Large-scale untargeted metabolomics experiments lead to detection of thousands of novel metabolic features as well as false positive artifacts. With the incorporation of pooled QC samples and corresponding bioinformatics algorithms, those measurement artifacts can be well quality controlled. However, it is impracticable for all the studies to apply such experimental design.

**Results:** We introduce a post-alignment quality control method called genuMet, which is solely based on injection order of biological samples to identify potential false metabolic features. In terms of the missing pattern of metabolic signals, genuMet can reach over 95% true negative rate and 85% true positive rate with suitable parameters, compared with the algorithm utilizing pooled QC samples. genuMet makes it possible for studies without pooled QC samples to reduce false metabolic signals and perform robust statistical analysis.

**Availability and implementation:** genuMet is implemented in a R package and available on https://github.com/liucaomics/genuMet under GPL-v2 license.

**Contact:** Liming Liang

**Supplementary information:** Supplementary data are available at ….

## 1 Introduction

Metabolomics consists of methods and techniques measuring the metabolite profile of biofluids (Alonso, et al., 2015). It has been successfully applied to both biomarker discovery and molecular mechanism inference (Johnson, et al., 2016). Untargeted metabolomics experiments give a holistic profiling of metabolites, including both known metabolites and structurally novel metabolites (Vinayavekhin and Saghatelian, 2010). However, due to technical variations like drift in chromatographic and mass spectrometric performance overtime (Dunn, et al., 2012) and miscalibration, false signals may be detected as untargeted metabolic features, which could reduce the power of subsequent statistical analysis and lead to unreliable biological causality inference.

Special experimental designs and innovative informatics tools can help eliminate measurement artifacts. Quality control (QC) samples like pooled QC samples and commercial standard biofluids are routinely included in large-scale metabolic studies mainly for three reasons: equilibrate analytical platforms after maintenance, calculate technical precision for quality assurance and integrate data from different analytical batches (Dunn, et al., 2011). Recently, a post alignment QC algorithm based on intermittent pooled QC samples called MetProc (Chaffin, et al., 2019), is developed to identify potential measurement artifacts.

However, pooled QC samples are not always available. In large-scale studies, sample collection may not be completed before sample preparation and analysis. In studies with limited sample availability, like bile, tears and interstitial fluid metabolic profiling, pooled sample preparation is very difficult as well (Dunn, et al., 2011). As it is impossible for commercial or synthetic biofluid to cover all the untargeted metabolites, they are not good surrogate for pooled QC samples. Therefore, new QC tools for detecting measurement artifacts that not relies on QC samples are in great need. Here, we present a R package genuMet which is solely based on the missing pattern of metabolic features along the injection order to separate genuine untargeted metabolic features from potential measurement artifacts without any QC samples. We compare its performance with MetProc using metabolic data from the PREDIMED (PREvención con DI-eta MEDiterránea) study (www.predimed.es).

## 2 Description

We focus on the missing rate pattern of metabolic signal along the injection order instead of the absolute signal intensity. A sliding window method is employed in genuMet. The main framework is illustrated in Figure 1. For each metabolic feature, a window will slide across the injection order, missing rate is calculated within each window, and each metabolite will end up with a series of missing rates. As we assume that the injection order of all the samples has been randomized, any metabolic feature with structured missing rate pattern tend to be measurement artifacts. Four metrics are employed to characterize the missing pattern: variance of missing rates, number of switches, the length of longest block and mean missing rate (see supplement information for details).

**Figure 1.**
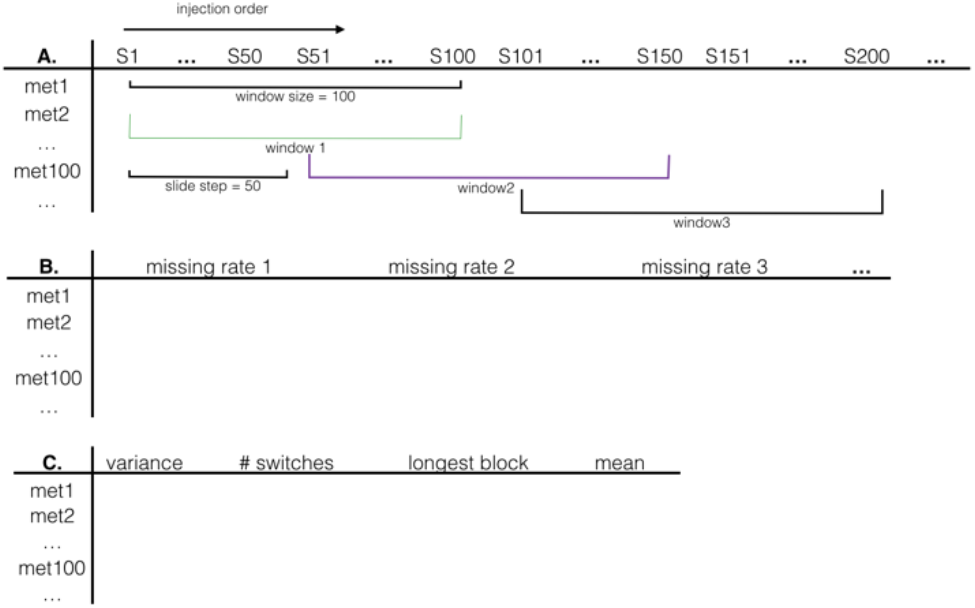
(A) The raw metabolic signal matrix. The columns are biological samples in injection order and rows are metabolic features. A window of window size 100 slides across the injection order with slide step 50. (B) Missing rate matrix. (C) Summary statistics matrix.

The four metrics are then combined to decide whether a metabolic feature is a potential measurement artifact or not. One possible criterion is that metabolic features with large missing rate variance (default >= 0.03) or high mean missing rate (default >=0.95) are classified as measurement artifact, and for those with small variance, if they have large number of switches (default >=1) and short longest block (default<=35), it should be classified as measurement artifact as well.

## 3 Results and discussion

We take the predicted result of MetProc as gold standard and assess the performance of genuMet with lipid metabolites generated for the PREDIMED study (Guasch-Ferre, et al., 2016; Martinez-Gonzalez, et al., 2012). This data set consists of 1,989 biological samples and 6,359 metabolic features. Suppl Fig. 1 and Suppl Fig. 2 indicate the individual prediction ability of the four metrics. With default parameters, our method identifies 1279 artifacts, reaching over 95% true negative rate and 85% true positive rate. Moreover, in some special scenarios, genuMet could even detect artifacts that are neglected by MetProc (Suppl Fig. 3A & 3B). However, without the information of pool QC sample, genuMet lacks the ability to characterize the local missing trends and detect rare metabolic features (Suppl Fig. 3C & 3D). The heatmaps (Suppl Fig. 4) suggests that genuMet does successfully separate true signals with random missing data and potential artifacts with structured missing pattern.

The comparison of genuMet with MetProc on PREDIMED metabolomics data demonstrates its ability to separate metabolic feature with structured missing pattern, which should be removed from downstream analysis. With suitable parameters, genuMet can give acceptable accuracy for identifying measurement artifact after metabolic feature detection and alignment. However, due to the lack of pooled QC samples, it is difficult to determine whether the local variation of missing rate should be attributed to technical effect or biological signal variation. The default parameters for separating metabolites were selected based on PREDIMED study, genuMet package provides flexible functions that can be adjusted to cater for the user’s need. Our method also provides a variety of graphical tools for quality control and visualization of the missing rate pattern. In summary, genuMet is the first post-alignment tool that allows large scale untargeted metabolomics studies without QC samples to distinguish potential genuine metabolic features from artifacts, which is of great significance for reproducibility and robust biological causality inference.

## Acknowledgements

The authors thank all the participants for their collaboration, all the PREDIMED personnel for their assistance and all the personnel of affiliated primary care centers for making the study possible.

## Funding

This work was supported by NIH research grant HL118264. The PREDIMED trial was supported by the official funding agency for biomedical research of the Spanish government, Instituto de Salud Carlos III (ISCIII), through grants provided to research networks specifically developed for the trial (RTIC G03/140, to Ramón Estruch, 2003-2005 and RTIC RD 06/0045, to Miguel A. Martínez-González, 2006-2013, and by Centro de Investigación Biomédica en Red de Fisiopatología de la Obesidad y Nutrición [CIBEROBN] and grants from Centro Nacional de Investigaciones Cardiovasculares (CNIC 06/2007), Fondo de Investigación Sanitaria–Fondo Europeo de Desarrollo Regional (PI04–2239, PI 05/2584, CP06/00100, PI07/0240, PI07/1138, PI07/0954, PI 07/0473, PI10/01407, PI10/02658, PI11/01647, P11/02505, PI13/00462 and PI13/01090), Ministerio de Ciencia e Innovación (AGL-2009–13906-C02 and AGL2010–22319-C03), Fundación Mapfre 2010, Consejería de Salud de la Junta de Andalucía (PI0105/2007), Public Health Division of the Department of Health of the Autonomous Government of Catalonia, Generalitat Valenciana (ACOMP06109, GVA-COMP2010–181, GVACOMP2011–151, CS2010-AP-111, and CS2011-AP-042), and Regional Government of Navarra (P27/2011).

### Conflict of Interest

none declared.

## Notes

https://github.com/liucaomics/genuMet

