## Supplemental Figure 1-4 for "genuMet: distinguish genuine untargeted metabolic features without quality control samples"

### Supplementary Information

1. Description of the four metrics
2. Supplementary Figure 1. Distribution of four genuMet metrics
3. Supplementary Figure 2. ROC curve for individual metric.
4. Supplementary Figure 3. Missing rate series of four example metabolic features
5. Supplementary Figure 4. Heatmap of missing status of metabolic features

### genuMet: distinguish genuine untargeted metabolic features without quality control samples

Cao L[1,15], Clish C[2], Hu FB[1], Martínez-González MA[1,3,4], Razquin C[3,4], Bullo-Bonet M[4,5], Corella D[4,6], Gómez-Gracia E[4,7], Fiol M[5,8], Estruch R[4,9], Lapetra J[4,10], Fitó M[4,11], Arós F[4,12], Serra-Majem L[4,13], Ros E[3,14], and Liang L[1,*]

[1]Harvard TH Chan School of Public Health, Boston, MA, USA, [2]Broad Institute of MIT and Harvard, Cambridge, MA, USA, [3]University of Navarra, Pamplona, Spain, [4]Instituto de Salud Carlos III (ISCIII), Madrid, Spain, [5]Universitat Rovira i Virgili, Reus, Spain, [6]University of Valencia, Valencia, Spain, [7]University of Malaga, Malaga, Spain, [8]Instituto de Investigación Sanitaria de Palma, Palma de Mallorca, Spain, [9]Institut d'Investigacions Biomèdiques August Pi i Sunyer, Barcelona, Spain, [10]San Pablo Health Center, Sevilla, Spain, [11]Parc de Salut Mar, Barcelona, Spain, [12]University Hospital of Alava, Vitoria, Spain, [13]University of Las Palmas de Gran Canaria, Las Palmas, Spain, [14]Institut d'Investigacions Biomèdiques August Pi i Sunyer, Barcelona, Spain, [15]Carnegie Mellon University, Pittsburgh, PA, USA

*To whom correspondence should be addressed.

##### 1. Description of the four metrics

Variance of missing rates for measurement artifacts tend to be large as these signals originate from unstable experiment shift, while the variance for genuine metabolite should be small, because after randomization the missing rate in each window tend to be similar (Supp Figure 1A)

A switch is defined as great contrast of missing rate between adjacent windows. When the threshold of missing rate contrast is large enough, very few switches will be detected in genuine metabolic feature. But when the threshold becomes small, some genuine metabolic feature may have extremely large number of switches (Supp Figure 1B).

A block is defined as the consecutive windows between two switches. The length of longest block of most true metabolic features equals to the number of windows as no switches are identified at all (Supp Figure 1C).

Measurement artifact generally has very high missing rate. Consequently, some obvious false signal may have small variance, small number of switches and full longest block, which is hard to be detected by the first 3 metrics. Mean missing rate can well measure the overall high missing rate of false signal. (Supp Figure 1D).

#### 2. Supplementary Figure 1

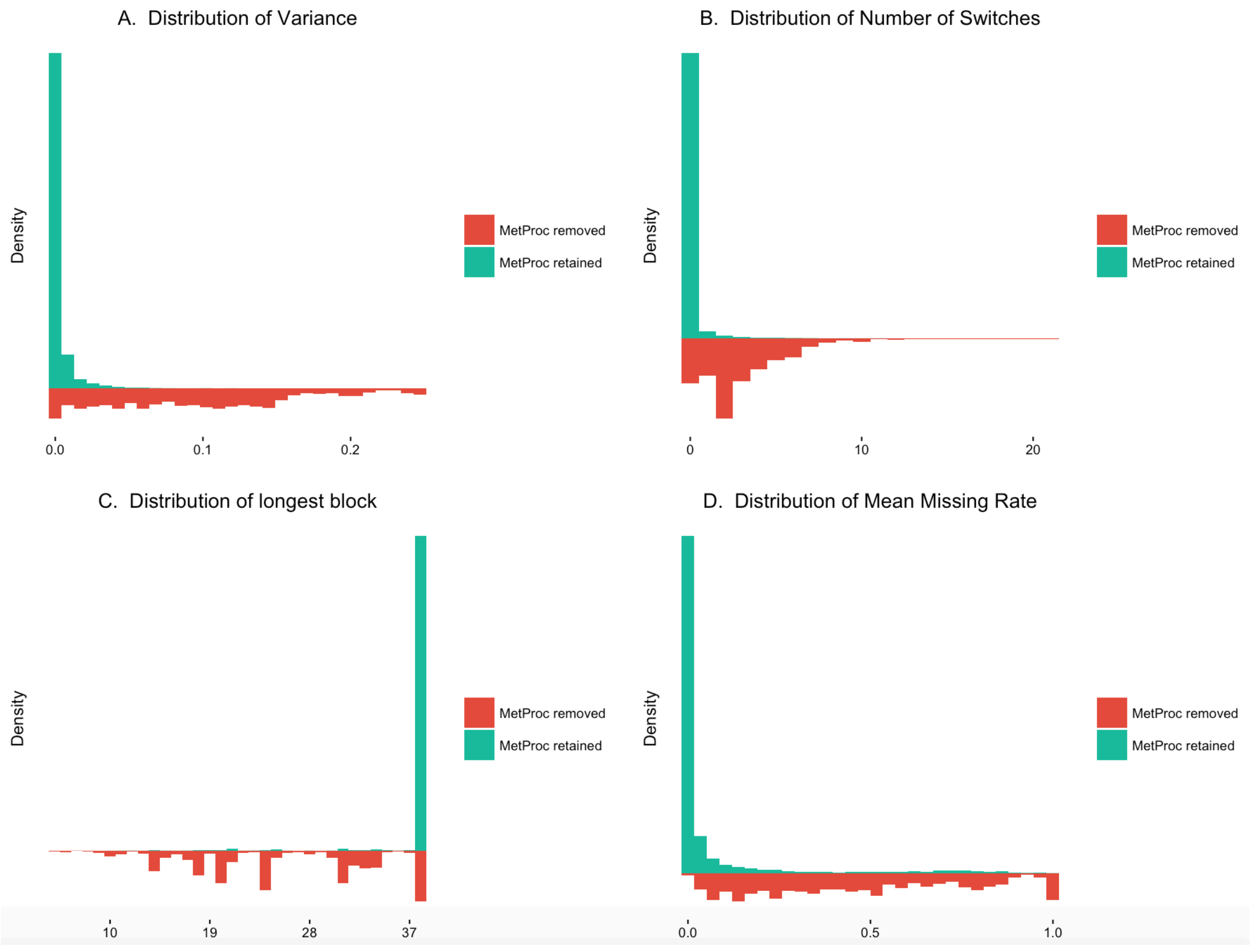


**Figure 1.** The distributions of the four metrics for removed and retained metabolic features predicted by MetProc. (a) Distribution of variance of missing rates. (b) Distribution of number of switches. (c) Distribution of length of longest block. (d) Distribution of mean missing rate.

**3. Supplementary Figure 2**

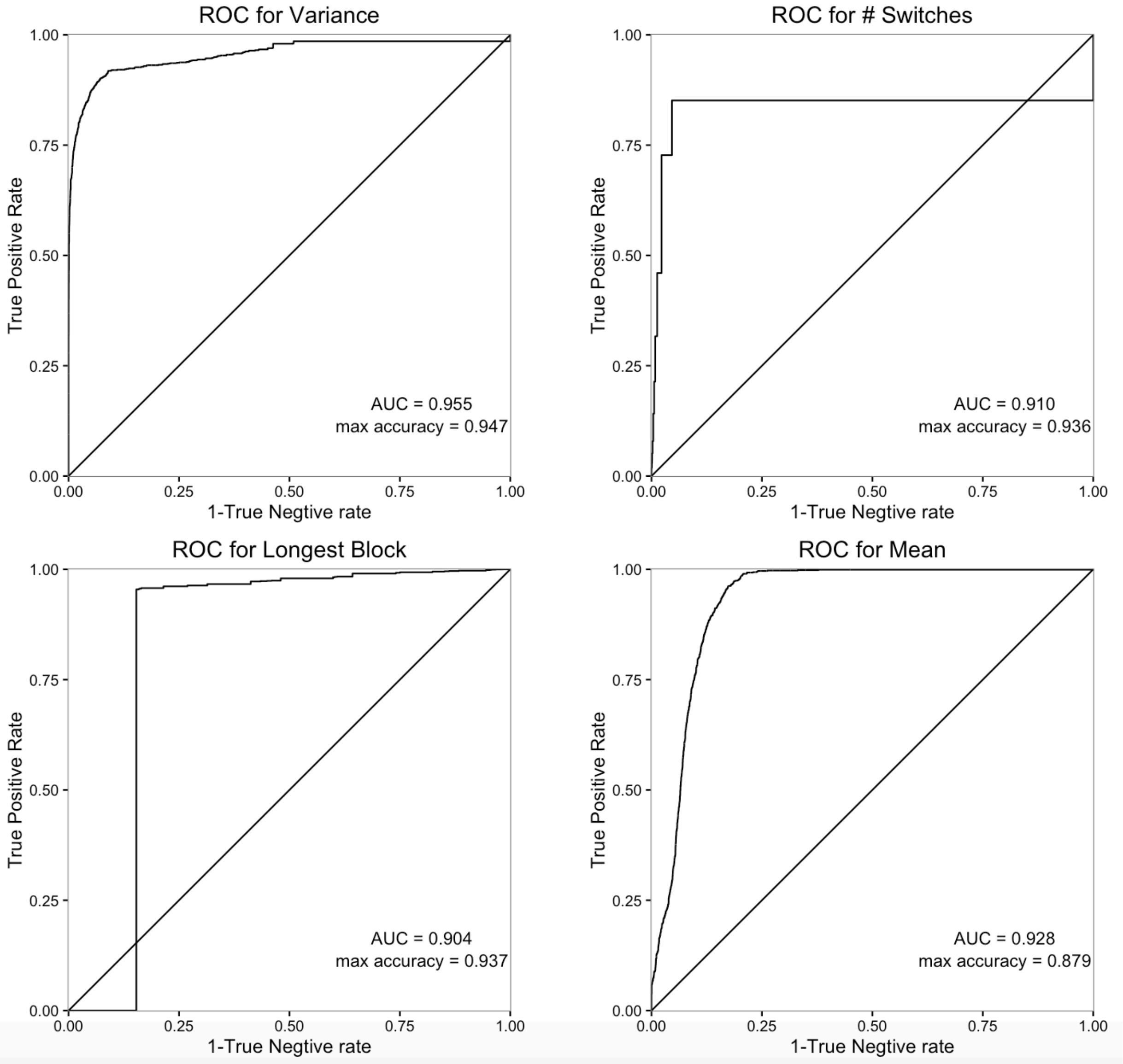


**Figure 2.** ROC curve for individual metric. The removed metabolic features predicted by MetProc are positives and retained are negatives.

**4. Supplementary Figure 3**

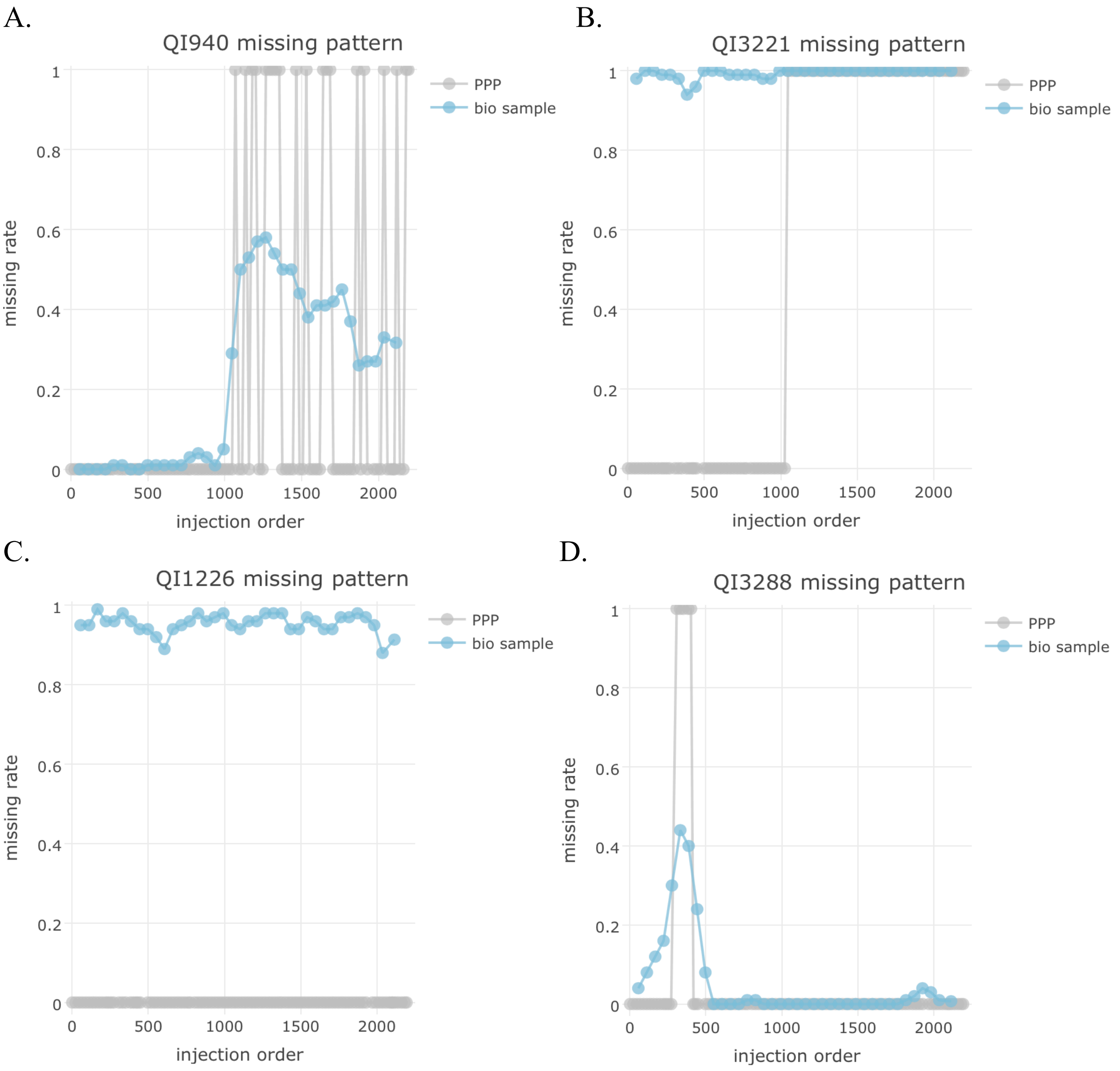


**Figure 3.** Missing pattern of four metabolic features.

(A) The missing rate series of QI940 shows that there is a huge missing rate change for both biological samples (blue) and pooled plasma (grey) due to the batch effects before and after the change of chromatography column around injection order 1000. QI940 should be classified as an artifact or low quality metabolic features. However, due to low PPP missing rate (Chaffin MD et al., 2016) and low correlation between PPP missing status and sample missing rate, MetProc finally fail to classify QI940 as problematic signal. Whereas, genuMet successfully identify this artifact with the metrics of variance of missing rate.

(B) A more extreme example is QI3221. It seems that after the change of column the missing rate of both pooled plasmas and biological samples go to 1 and this metabolic feature completely disappear. Undoubtedly, QI3221 is problematic and should be removed. But Metproc retain this feature due to low correlation between PPP status and biological sample missing rate.

(C) Without the PPP missing status, genuMet also fails to detect genuine but rare metabolic features. As QI1226 is never missing in the pooled plasmas, we can safely conclude that it should be a genuine metabolic feature. However, its biological missing rate remains very high in all analytical batches, indicating it only appears in a small portion of samples. To decrease false negative detection, genuMet naively remove metabolic features with high missing rate in biological samples and identified QI1226 as measurement artifact.

(D) Without the information of pool QC sample, genuMet lacks the ability to characterize the local missing trends. For ex- ample, the missing rate for QI3288 of both pooled plasma and biological samples stays high before the first 500 samples, indicating that the missing is caused by technical effect of first several analytical batches and it should be identified as problematic metabolic feature. However, due to the lack of the PPP missing status, genuMet can only rely missing pattern of biological samples, which makes it hard to distinguish local technical variation from biological variation and leads to false negative detection.

#### 5. Supplementary Figure 4

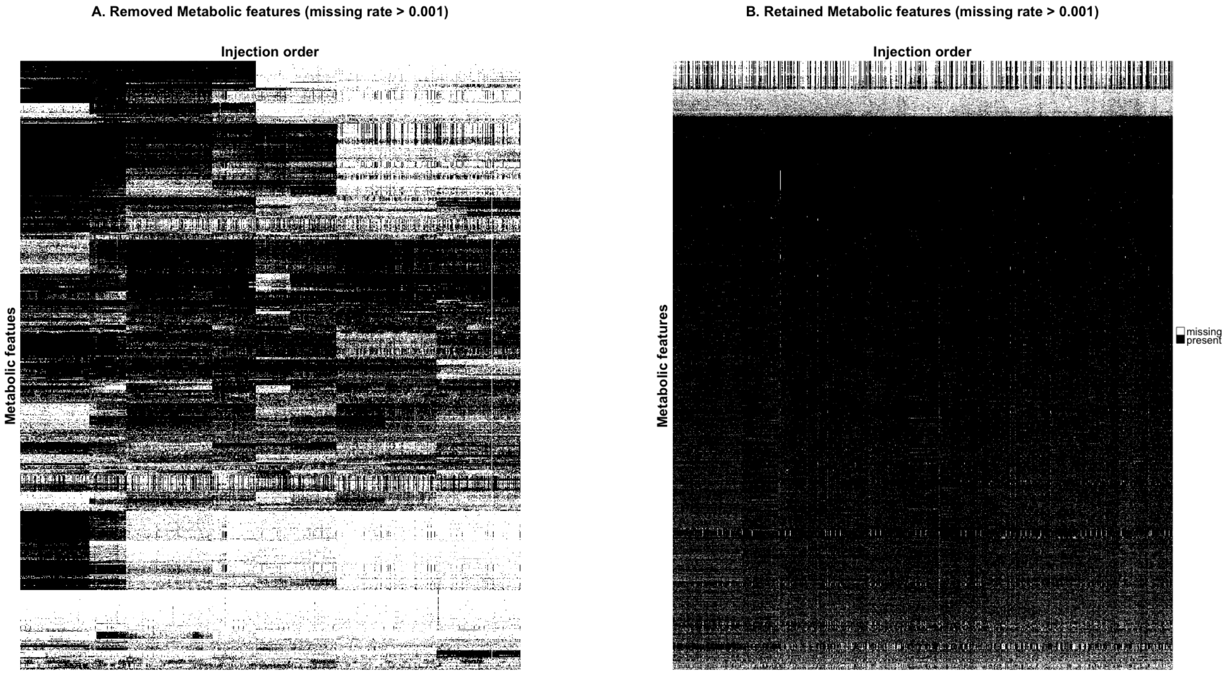


**Figure 4.** Heat map of missing status for (A) removed and (B) retained metabolic features of genuMet.
